# Crown protein is dispensable for OmRV entry but required for efficient transmission

**DOI:** 10.64898/2025.12.03.692111

**Authors:** Gabriela Garcia Hernández, Han Wang, Clara Manesco, Kenta Okamoto

## Abstract

Non-enveloped icosahedral dsRNA viruses, such as taxons of the Totiviridae family, have evolved surface features that enable infection of multicellular hosts through receptor binding and membrane penetration via the canonical endosomal pathway. Unlike Totiviruses that infect unicellular hosts, toti-like viruses, including artiviruses, infect invertebrate multicellular hosts and possess an additional surface crown protein (CrP) on their conserved T=1 icosahedral capsid. The mosquito-specific artivirus Omono River virus (OmRV) also expresses this CrP, which has been suggested to facilitate viral infection and propagation, although it may not be essential. To clarify its role in viral infection, we generated a CrP deletion mutant of OmRV to directly assess its functional significance. The absence of CrP does not affect infectivity in mosquito cells or alter the capsid structure. However, CPE and viral propagation are still affected. These findings indicate that CrP is dispensable for OmRV infection but may play a crucial role in later stages of cell-to-cell transmission, providing important insight into its previously unclear function in the viral life cycle.

## Introduction

Totiviridae is a large family of double-stranded RNA (dsRNA) viruses that primarily infect unicellular hosts such as yeast and protozoan species (Tighe et al., 2022). More recently, toti-like viruses have been discovered in multicellular hosts, including arthropods such as mosquitoes and shrimp, as well as in fish (Haugland et al., 2011; Isawa et al., 2011; Poulos et al., 2006; Tighe et al., 2022). These newly identified viruses have been reclassified into distinct taxonomic families: Artiviridae (arthropod toti-like viruses) (Zhai et al., 2010), and Pistolviridae (piscine toti-like viruses) (Sandlund et al., 2021). Several more taxonomic genera are also suggested including Tricladivirus (Triclad flatworm toti-like virus) (Burrows et al., 2020; Tighe et al., 2022). One such virus, Omono River virus (OmRV), was isolated from *Culex* mosquitoes and is classified under the family Artiviridae (Isawa et al., 2011). Importantly, some of these toti-like viruses are significant pathogens in aquaculture, including Infectious Myonecrosis Virus (IMNV), which affects shrimp (Poulos et al., 2006), and Piscine Myocarditis Virus (PMCV), which infects salmonid fish (Haugland et al., 2011). These viruses raise both economic concerns for the fisheries industry and environmental concerns for wild shrimp and fish populations (Prasad et al., 2017; Svendsen et al., 2019).

The most critical distinction between Totiviridae and Artiviridae/Pistolviridae lies in their transmission mechanisms. Totiviridae members, such as *Saccharomyces virus L-A* (ScV-L-A) infecting yeast and *Trichomonas vaginalis virus 1* (TVV1) infecting protozoa, rely on intracellular transmission. They are passed vertically through host cell division and mating, common in unicellular organisms (Ghabrial, 1998; Goodman et al., 2011; Ihrmark et al., 2002). In contrast, Artiviridae and Pistolviridae utilize extracellular transmission, involving viral entry into host cells and exit via budding. A particular exception is Giardia lamblia virus (GLV), which infects the unicellular protozoan *Giardia lamblia* belonging to Totiviridae, however is phylogenetically close to Artiviridae (Isawa et al., 2011; Tighe et al., 2022). GLV employs an extracellular transmission mechanism similar to Artiviridae viruses (Marucci et al., 2021; Tai et al., 1993). It is a long-standing hypothesis that this extracellular transmission requires the acquisition of surface structural features that enable the virus to bind host cell receptors and penetrate endosomal membranes, while these features are not required for non-enveloped dsRNA viruses transmitted solely via intracellular mechanisms (Nibert and Takagi, 2013).

Both Totiviridae and Artiviridae viruses express a capsid protein (CP) and an RNA-dependent RNA polymerase (RdRp) to form a T=1 icosahedral capsid lattice. In OmRV and IMNV, the CP is further processed to generate a major capsid protein (MCP) accompanied by several smaller proteins (P1-4) (Isawa et al., 2011; Shao et al., 2021). One of these smaller proteins is the crown protein (CrP), as previously also described as surface fiber or protrusion, which is structurally located at the 5-fold vertices of the icosahedral capsid (Shao et al., 2021; Tang et al., 2008). Although the OmRV LZ strain displays CrPs on the capsid surface, the OmRV AK4 strain does not, despite both strains expressing CrPs (Okamoto et al., 2020, 2016; Shao et al., 2021). Similar to the OmRV AK4 strain, GLV neither expresses CrPs nor presents them on the capsid surface (Wang et al., 2024). However, detailed structural comparisons between Totiviridae and Artiviridae viruses reveal that both OmRV and GLV have acquired several loop structures on the capsid surface that are absent in Totiviridae viruses relying on intracellular transmission (Okamoto et al., 2020; Wang et al., 2024). Although CrP and the acquired surface loops are thought to be important for extracellular transmission, their function remains unclear. OmRV propagation is inhibited by CrP-specific antibodies (Shao et al., 2021) or by a point mutation in OmRV MCP that weakens the MCP-CrP interaction (Wang et al., 2022), suggesting that CrP promotes infection, but is presumably not essential. In pistovirus PMCV, a different but additional protein is expressed in the ORF3, however the function is so far unknown (Haugland et al., 2011).

Here, we have generated a CrP-deletion mutant and characterized its transmission and structure to elucidate the role of CrP in OmRV infection, aiming to deepen our understanding of the still unclear extracellular transmission mechanism of Artiviridae viruses.

## Materials and Methods

### Generation of OmRV CrP deletion infectious DNA clone

The plasmid OmRV-WT/pACYC177 was constructed in a previous study (Wang et al., 2022). The strategy for generating OmRV-CrPdel is shown in Fig. 1. Fragment 1 encodes the genes of the putative proteins P1 to P4 and part of the MCP within ORF1. The deletion spans nucleotides 748 to 1,113 bp, corresponding to the N-terminal region of P3, thereby removing the entire CrP gene coding region except for the proteolytic cleavage sites. The designed plasmid includes an upstream SP6 promoter sequence for in vitro transcription. From the designing, CrP-gene deletion Fragment 1 (2,491 bp) was synthesized in a pMA plasmid (OmRV-CrPdel-fragment1/pMA) using the GeneArt Custom Gene Synthesis service (Thermo Fisher Scientific). The remaining part of the OmRV genome, Fragment 2 (4,748 bp), was inserted into the pACYC177 plasmid (OmRV-fragment2/pACYC177). Fragment 1 was subsequently cloned into OmRV-fragment2/pACYC177 to assemble a full-length infectious DNA clone of OmRV-CrPdel (OmRV-CrPdel/pACYC177). The complete DNA sequence of OmRV-CrPdel/pACYC177 was confirmed by Sanger sequencing.

**Fig. 1.**
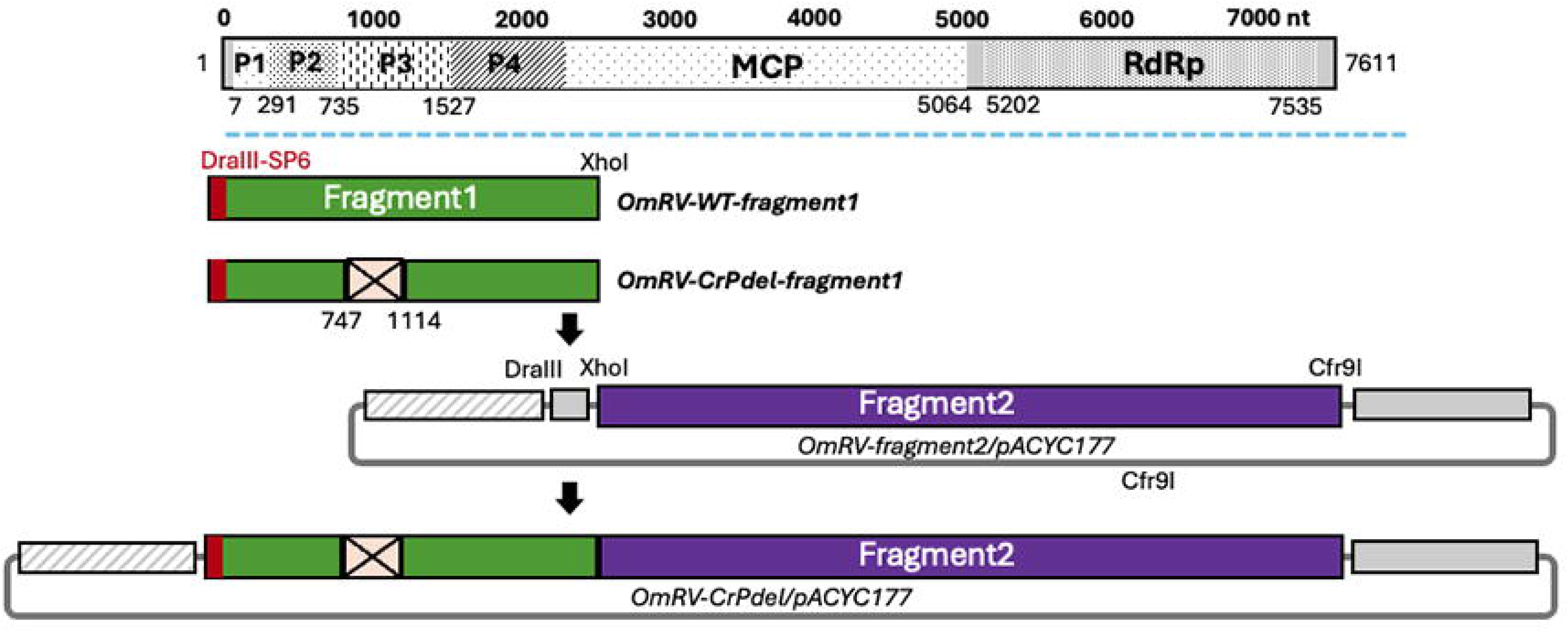
Design and generation of the OmRV-CrPdel/pACYC177 plasmid. The OmRV-CrPdel-fragment1 was designed to delete nucleotides 748–1113, corresponding to the N-terminal region of the P3 protein. This CrP-deleted fragment (OmRV-CrPdel-fragment1) was cloned into the DraIII and XhoI sites of OmRV-fragment2/pACYC177. The resulting plasmid, OmRV-CrPdel/pACYC177, contains the complete genome sequence of the OmRV-AK4 strain, except for the deleted region.

### In vitro generation of OmRV-WT and OmRV-CrPdel mutant

Mosquito C6/36 cells were cultured in Minimal Essential Medium (MEM) supplemented with 10% (v/v) fetal bovine serum (Sigma-Aldrich, F9665), 2 mM L-glutamine, non-essential amino acids, and penicillin–streptomycin at 28°C in a 5% CO₂ atmosphere. These cells were used for generating recombinant OmRV.

Both OmRV-WT/pACYC177 and OmRV-CrPdel/pACYC177 plasmids were linearized at the Cfr9I restriction site, purified, and used as templates for in vitro transcription to synthesize full-length infectious viral RNAs. In vitro transcription was performed using 5 µg of each linearized plasmid with the mMESSAGE mMACHINE High-Yield Capped RNA Transcription SP6 Kit (Ambion, AM1340). To optimize transcript yield, 6 µL of 20 mM GTP was added to the reaction mixture. The resulting viral RNAs were purified and resuspended in RNase-free water, then stored at −80°C until transfection.

For transfection, 2.5 µg of viral RNA was introduced into C6/36 cells cultured in 6-well plates using Lipofectamine 2000 reagent (Thermo Fisher Scientific, 11668027). The infected cell fluid (ICF) was collected for the further assays.

### Propagation kinetics and Infectivity assay

For propagation assay, after viral RNA transfection or virus inoculation at a multiplicity of infection (MOI) of 10, 220 µL of culture supernatant was collected at 0 (2 h), 1, 2, 3, 4 and 5 days post-infection (dpi). Each collected sample was centrifuged to separate the ICF from cell debris. The viral titers in the ICF were quantified using reverse transcription quantitative PCR (RT-qPCR). For the infectivity assay, C6/36 cells were inoculated with either the WT or CrPdel mutant virus at an MOI of 10. The cells were incubated for 2 h on ice, followed by 2 h at 28°C. After incubation, the ICF was removed, and the cells were gently rinsed with PBS to eliminate unbound viral particles. The cells were then lysed, and total RNA was extracted using TRIzol reagent. A mock-infected control was included as a negative control, in which nuclease-free water was used instead of viral RNAs or viral particles. Intracellular viral RNA levels were quantified by RT-qPCR.

### RT-qPCR titration

To determine the presence of viruses in the ICF, viral dsRNA was extracted using the PureLink Viral RNA/DNA Mini Kit (Thermo Fisher Scientific, 12280050). The PCR primers (OmRV-AK4-519F: 5’-TGTGTATAAGGTTGGGTCGGAAG-3’ and OmRV-AK4-674R: 5’-GACAACAAACACATAGGACAGAA-3’) and a probe with FAM fluorophore at the 5’-end and BHQ1 quencher at the 3’-end (OmRV-AK4-Probe: 5’-FAM-TATAACCAAGCCTTTGCTGGGCGT-BHQ1-3’) were used. In a 35 µl reaction mix, AgPath-ID One-Step RT-PCR (Applied Biosystems, AM1005) enzyme mix and the buffer were mixed with 4 µl of a RNA sample. Each PCR primer had a final concentration of 400 nM and the probe had a final concentration of 120 nM. Reverse transcription was carried out at 45°C for 10 min, followed by enzyme inactivation at 95°C for 10 min. Amplification consisted of 40 cycles of 95°C for 15 s and 60°C for 45 s. All reactions were performed on a QuantStudio 6 Flex System (Applied Biosystems). Two technical replicates were included to confirm reproducibility. The number of viral particles in each sample was calculated using a standard curve generated from a plasmid sample of known concentration, as described previously (Wang et al., 2022).

### Cryo-EM structural determination

OmRV-WT and OmRV-CrPdel were propagated in confluent C6/36 cells grown in ten 175 cm² flasks. Purification of the recombinant OmRV followed a previously described procedure (Okamoto et al., 2020). The final concentration of purified viruses ranged from 1 to 3 mg/mL and was used for cryo-EM grid preparation. Cryo-EM grids were prepared by applying 3 µL of purified sample onto glow-discharged holey carbon grids (Quantifoil R1.2/1.3, Cu 300 mesh) and flash-freezing them in liquid ethane after blotting for 3 s at 4 °C and 100% humidity using a Mark IV Vitrobot (Thermo Fisher Scientific).

Data collection was performed using a Glacios cryo-EM (Thermo Fisher Scientific) equipped with a Falcon 3 imaging sensor at the Uppsala Cryo-EM facility. Image processing and 3D reconstruction were carried out using CryoSPARC version 4.3.1 (Punjani et al., 2017). Image movies were acquired with a pixel size of 0.952 Å/pixel, a total electron exposure of 29.57 e⁻/Å², and defocus values ranging from 0.7 to 1.5 µm underfocus. A total of 17,916 and 1,323 movies were collected for OmRV-WT and OmRV-CrPdel, respectively. After motion correction and contrast transfer function (CTF) estimation, 31,506 particles of OmRV-WT and 12,488 particles of OmRV-CrPdel were extracted and used for 3D reconstruction imposing icosahedral symmetry. The final 3D reconstructions reached resolutions of 3.1 Å and 2.9 Å for OmRV-WT and OmRV-CrPdel, based on the Fourier Shell Correlation (FSC) 0.143 cutoff. Atomic models of recombinant OmRV-WT and OmRV-CrPdel MCPs were refined against the obtained maps and the previously deposited OmRV structure (PDB ID: 6S2C) using Coot version 1.0.06 (Emsley et al., 2010) and PHENIX version 1.21.2 (Liebschner et al., 2019). Model validation statistics are summarized in Supplementary Table S1. Cryo-EM maps and atomic models were rendered using UCSF ChimeraX (Meng et al., 2023).

## Results and Discussion

### Propagation and infectivity of OmRV-WT and OmRV-CrPdel

The propagation of OmRV-WT and OmRV-CrPdel was initially compared following lipofection with equal amounts of viral RNA (Supplementary Fig. S1). No discernible differences in cytopathic effect (CPE) were observed between the two viruses from 1 to 5 days post-infection (dpi) (Supplementary Fig. S1A). However, across several independent experiments, higher viral RNA levels of OmRV-CrPdel were observed in ICF at both 0 and 1 dpi (Supplementary Fig. S1B). This is most likely because the amount of viral RNA introduced into the cells differed between OmRV-WT and OmRV-CrPdel, owing to variations in lipofection efficiency related to their genome sizes, even though the same nominal amount was used. Despite these differences, the fold changes in viral RNA levels were comparable between the two viruses (Supplementary Fig. S2C). This discrepancy in the initial RNA load complicated direct comparisons of CPE and propagation efficiency. To more accurately evaluate CPE, propagation efficiency and infectivity, subsequent analyses were conducted using purified OmRV-WT and OmRV-CrPdel preparations standardized to identical infectious titers.

The observed CPE differed markedly between cells infected with OmRV-WT and those infected with OmRV-CrPdel (Fig. 2A). The regular C6/36 cells exhibit an adherent and measure approximately 5-10 μm in diameter. The typical CPE caused by OmRV infection is characterized by cell rounding, enlargement, and detachment (Isawa et al., 2011; Wang et al., 2022), as seen in OmRV-WT-infected cells at 3 and 5 dpi (Fig. 2A). In contrast, cells infected with OmRV-CrPdel displayed many considerably enlarged, amorphous cells (> 100 μm in diameter) at 3 dpi, likely resulting from cell fusion (Fig. 2A). By 5 dpi, these enlarged cells had developed into large, round cells (Fig. 2A). Although CPE appears more severe in OmRV-CrPdel-infected cells at 3 dpi, the ICF viral titer is substantially lower than that of OmRV-WT (Fig. 2B). However, by 5 dpi, the OmRV-CrPdel titer reaches levels comparable to those of OmRV-WT (Fig. 2B). Despite the differences in CPE and propagation efficiency, the intracellular viral titers measured 2 h post-inoculation are unaffected by the absence of CrP (Fig. 2C).

**Fig. 2.**
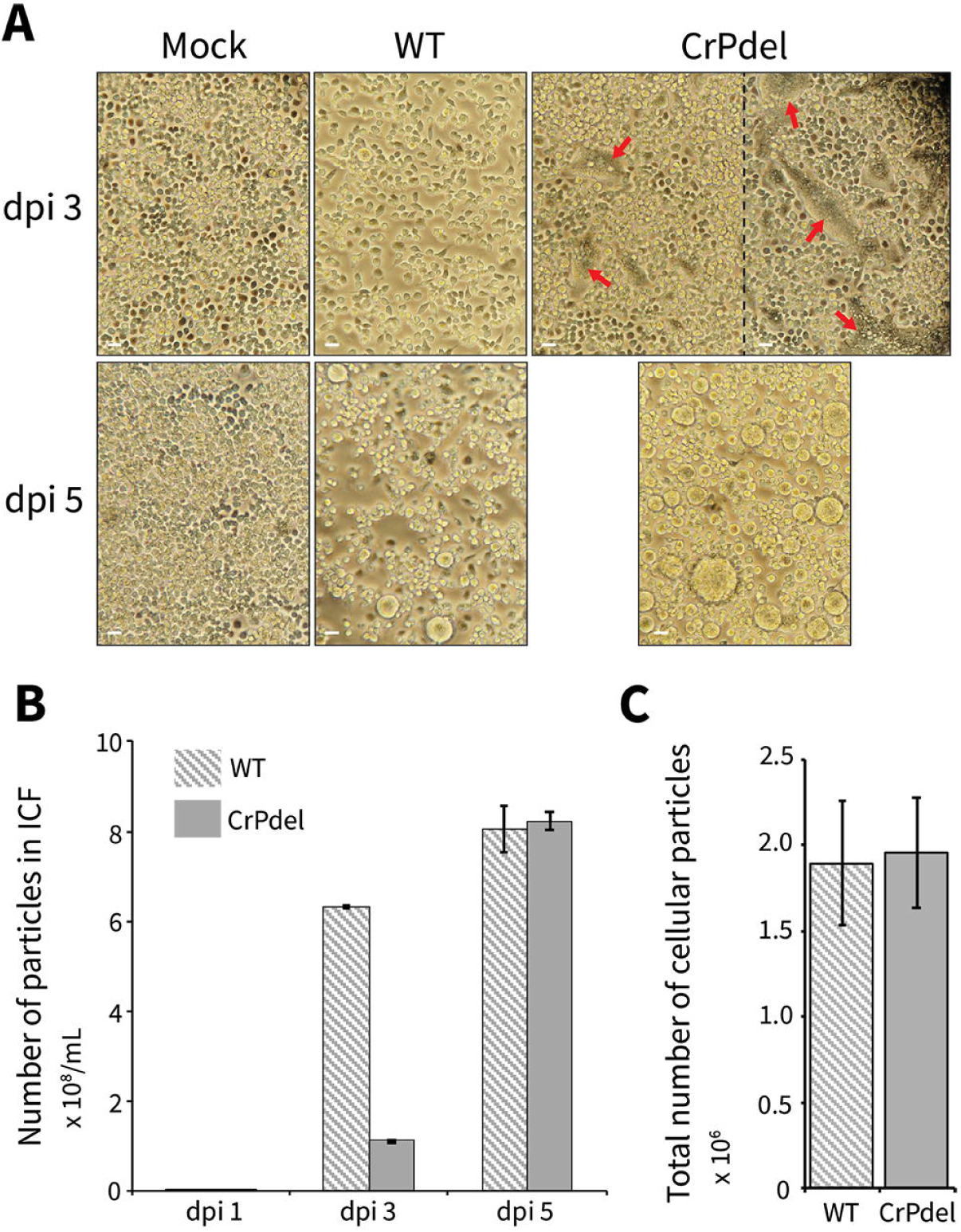
Propagation curve and infectivity of OmRV-WT and OmRV-CrPdel. **(A)** CPE in C6/36 cells at 3 and 5 dpi. C6/36 cells were infected with OmRV-WT or OmRV-CrPdel at an MOI of 10. Mock indicates uninfected control cells. The scale bars indicate 50 μm. **(B)** Propagation curves of OmRV-WT and OmRV-CrPdel. Cells were infected at an MOI of 10. The result is based on three technical replicates. **(C)** Intracellular virus titers measured 2 h post-inoculation (MOI = 10) for OmRV-WT and OmRV-CrPdel. Virus titers were determined from three biological replicates. The P value between OmRV-WT and OmRV-CrPdel is 0.916 (p > 0.05), based on three biological replicates.

It is unexpected that CrP does not affect the early stages of infection during the viral multiplication cycle. These results contrast with previous studies. However, earlier mutagenesis work introduced a point mutation on the capsid surface rather than directly on CrP to weaken the CrP-MCP interaction (Wang et al., 2022). Such mutations likely altered the surface loop structures of the capsid, which are thought to be critical for viral infection (Okamoto et al., 2020; Wang et al., 2024). Another study used an anti-CrP antibody to inhibit infection (Shao et al., 2021); however, the relatively large size of the antibody could have indirectly influenced the infectivity of OmRV. In both cases, the enhancement of infectivity by CrP was not pronounced and appeared to be minor (Shao et al., 2021; Wang et al., 2022). In this study, it is directly demonstrated that CrP is not involved in infection during cell entry in C6/36 mosquito cells. CrP might participate in an alternative extracellular transmission pathway, such as virus budding from cells. Although the extracellular viral titer remains relatively low despite the presence of extremely enlarged cells in OmRV-CrPdel-infected cultures, this observation may indicate an impairment in viral release, causing virions to be retained within the cells for a longer period. Further molecular studies will be required to fully elucidate the functions of CrP. Nonetheless, its role should be re-examined, focusing on the later stages of viral transmission.

### Structural comparison between OmRV-WT and OmRV-CrPdel

Currently, two structures of OmRV, the AK4 and LZ strains from original isolates, have been determined (Okamoto et al., 2020, 2016; Shao et al., 2021). Several minor structural variations of the MCP are observed in the OmRV LZ strain structures with or without CrP, including a slight tilting of the MCP in the absence of CrP (Shao et al., 2021). In the OmRV AK4 strain, CrP is unlikely to localize on the capsid surface because of weak interactions, yet it could remain detectable in the purified virus fractions (Okamoto et al., 2016; Shao et al., 2021). Such global and local structural changes in the MCP induced by the presence of CrPs may influence infectivity as well as other yet-unidentified cellular functions. Therefore, to directly clarify these structural changes in the absence of CrPs in the OmRV AK4 strain, the structures of recombinant OmRV-WT and OmRV-CrPdel mutant viruses were determined at resolutions of 3.1 Å and 2.9 Å, respectively (Supplementary Fig. S2).

The raw cryo-EM particle images show approximately 50% genome-filled and 50% empty particles in both OmRV-WT and OmRV-CrPdel (Fig. 3A), consistent with observations from the original isolates (Okamoto et al., 2016). The overall structure of OmRV-CrPdel exhibits a T=1 icosahedral capsid with two MCP subunits in the asymmetric unit (Fig. 3B) and does not display any significant global structural differences (Supplementary Fig. S2B). The atomic models of the A and B subunits in OmRV-CrPdel fit well with those of OmRV-WT (Fig. 3C). Local structural variations (RMSD < 3 Å) are observed, particularly at the interface between the A and B subunits (Fig. 3D), although considering the resolutions of the compared models, these differences are not regarded as significant. Based on the OmRV-CrPdel structure, the presence or absence of CrP does not affect the MCP structure, including features previously observed (Shao et al., 2021) or those predicted to be involved in cell entry, such as acquired surface loops (Okamoto et al., 2020; Wang et al., 2024). These results suggest that CrP is not a critical factor of triggering structural changes in the capsid during viral cell entry, which is consistent with our infectivity assay (Fig. 2C).

**Fig. 3.**
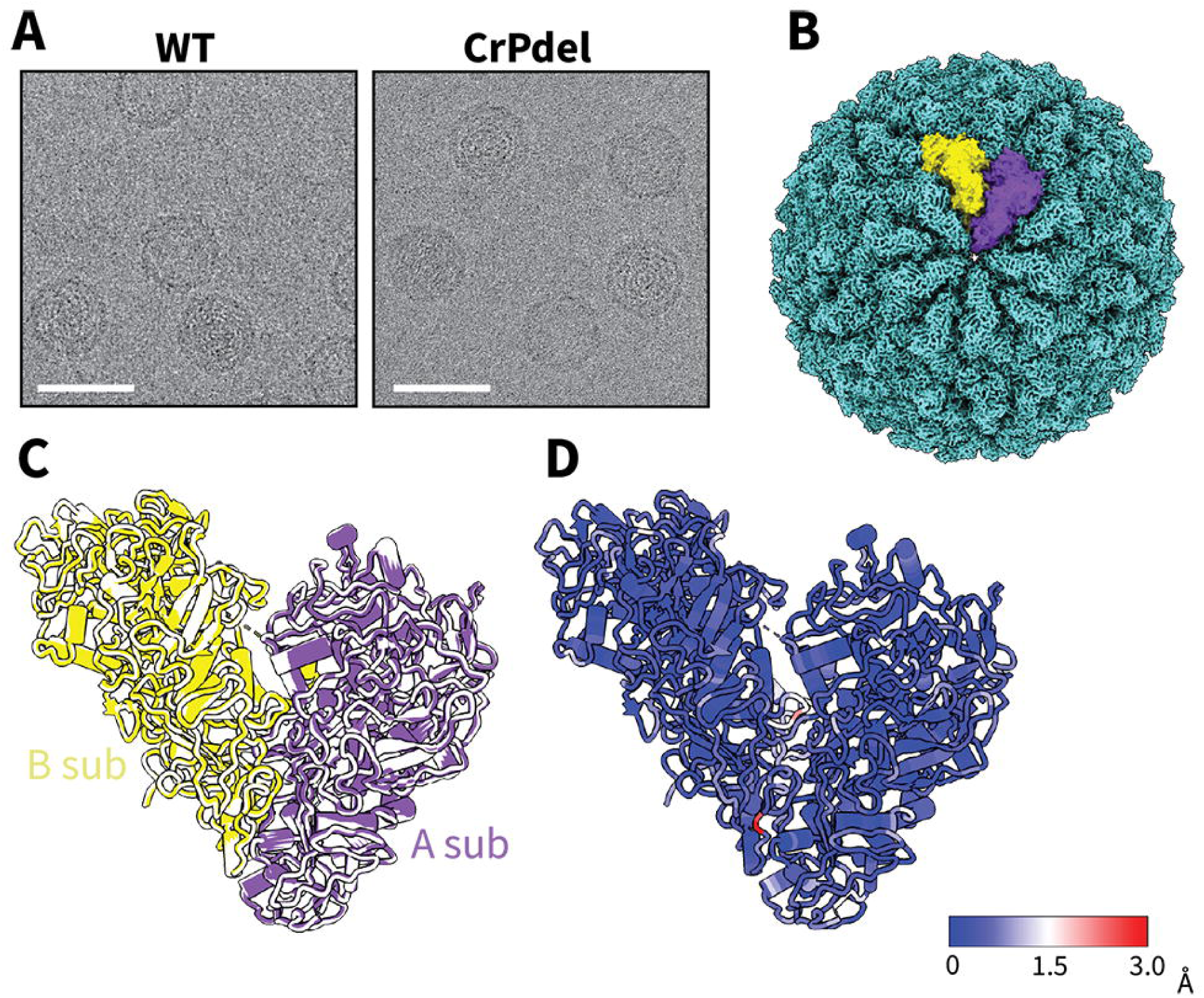
Structure of OmRV-WT and OmRV-CrPdel. **(A)** Raw particle images of OmRV-WT (left) and OmRV-CrPdel (right). Scale bar indicates 50 nm. **(B)** Icosahedral T=1 cryo-EM reconstruction of OmRV-CrPdel. Two subunits, A subunit (A sub) and B subunit (B sub), in an icosahedral asymmetric unit are colored in purple and yellow. **(C)** Superimposition of the atomic models of the A and B subunits in OmRV-WT and OmRV-CrPdel. **(D)** Structural variations between OmRV-WT and OmRV-CrPdel highlighted by residual root mean square deviation (RMSD) values.

Acquired surface structures or proteins in icosahedral dsRNA viruses infecting multicellular hosts are generally thought to play important roles in cell entry during extracellular transmission (Nibert and Takagi, 2013). However, there are two exceptions involving CrP. Megabirnavirus, which infects multicellular fungi, employs an intracellular transmission mechanism yet possesses a CrP-like protein on each 3-fold region (Wang et al., 2023). In contrast, GLV, which infects unicellular protozoa, utilizes an extracellular transmission mechanism but lacks CrPs (Wang et al., 2024). Together with the findings from OmRV-CrPdel, these observations strongly indicate that the extracellular cell entry mechanism of icosahedral dsRNA viruses, including artiviruses, does not depend on the CrP. However, we should not exclude that it may instead be related to in vivo aspects such as tissue tropism in viruses infecting multicellular hosts.

## Acknowledgement

C6/36 mosquito cells are kindly provided by Daisuke Kobayashi, Haruhiko Isawa, and Kyoko Sawabe, at National Institute of Infectious Diseases (NIID), Japan. We acknowledge the use of the Cryo-EM Uppsala facility for grid preparation, screening, and data collection, funded by the Department of Cell and Molecular Biology, the Disciplinary Domains of Science and Technology and of Medicine and Pharmacy at Uppsala University. Funding was provided by following agencies: Research Council of Norway (to Ø.E. and to K.O., grant number: 324266), the Swedish Research Council (to K.O., grant number: 2018-03387 and 2023-01857), and Carl Tryggers Stiftelse (to K.O., CTS 23:2703). For building the initial atomic model of OmRV-WT and OmRV-CrP, we thank Dags Macs, Babara Maria Nowak, and Yang Gao.

## Competing interests

The authors declare that they have no competing interests.

## Notes

### Competing Interest Statement

The authors have declared no competing interest.

