## Supplementary material for "Crown protein is dispensable for OmRV entry but required for efficient transmission": Suppl Table 1 and Figures 1 and 2

### Supplementary figures and table

| Atomic Models | PDB 9THH (OmRV-WT) | PDB 9THI (OmRV-CrPdel) |
|---|---|---|
| **Composition (#)** | | |
| Chains | 2 | 2 |
| Atoms | 13,177 | 13,177 |
| Residues | 1,727 | 1,727 |
| Water | 0 | 0 |
| Ligands | 0 | 0 |
| Bonds (RMSD) | | |
| Length (Å) (# > 4σ) | 0.003 (0) | 0.004 (0) |
| Angles (°) (# > 4σ) | 0.668 (4) | 0.883 (29) |
| MolProbity score | 2.29 | 2.52 |
| Clash score | 5.52 | 8.28 |
| **Ramachandran plot (%)** | | |
| Outliers | 0.58 | 0.76 |
| Allowed | 6.28 | 7.68 |
| Favored | 93.14 | 91.56 |
| **Rama-Z ( RMSD)** | | |
| whole | -0.71 (0.20) | -1.74 (0.20) |
| helix | 0.62 (0.24) | -1.06 (0.23) |
| sheet | -0.71 (0.39) | -0.94 (0.40) |
| loop | -0.93 (0.19) | -1.24 (0.19) |
| **Rotamer outliers (%)** | 5.33 | 5.61 |
| **Cβ outliers (%)** | 0.31 | 0.31 |
| **Peptide plane (%)** | | |
| Cis proline/general | 2.8/0.0 | 2.8/0.0 |
| Twisted proline/general | 0.9/0.0 | 0.9/0.0 |
| **CaBLAM outliers (%)** | 3.39 | 4.50 |
| **ADP (B-factors)** | | |
| Iso/Aniso (#) | 13,177/0 | 13,177/0 |
| min/max/mean | | |
| Protein | 28.67/194.73/70.25 | 13.27/128.59/46.33 |

| Cryo-EM maps | EMD-55927 (OmRV-WT) | EMD-55928 (OmRV-CrPdel) |
|---|---|---|
| **Box** | | |
| Lengths (Å) | 117.10,142.80,98.06 | 116.14,139.94,98.06 |
| Angles (°) | 90.00, 90.00, 90.00 | 90.00, 90.00, 90.00 |
| **Supplied Resolution (Å)** | 3.1 | 2.9 |
| **Resolution Estimates (Å)** | Masked (Unmasked) | Masked |
| d FSC (half maps; 0.143) | 2.9 | 2.9 |
| d 99 (full/half1/half2) | 26/5.4/5.4 | 2.4/3.6/3.6 |
| d model | 3.2 | 3.0 |
| d FSC model (0/0.143/0.5) | 2.9/3.1/3.3 | 2.3/2.8/3.1 |
| **Map min/max/mean** | -1.95/2.90/0.20 | -3.10/5.48/0.14 |

| Atomic models vs Cryo-EM maps | | |
|---|---|---|
| CC (mask) | 0.86 | 0.78 |
| CC (box) | 0.50 | 0.46 |
| CC (peaks) | 0.27 | 0.28 |
| CC (volume) | 0.84 | 0.73 |
| EMRinger | 2.72 | 2.16 |

**Supplementary Table S1 Validation statistics for cryo-EM and atomic models.**

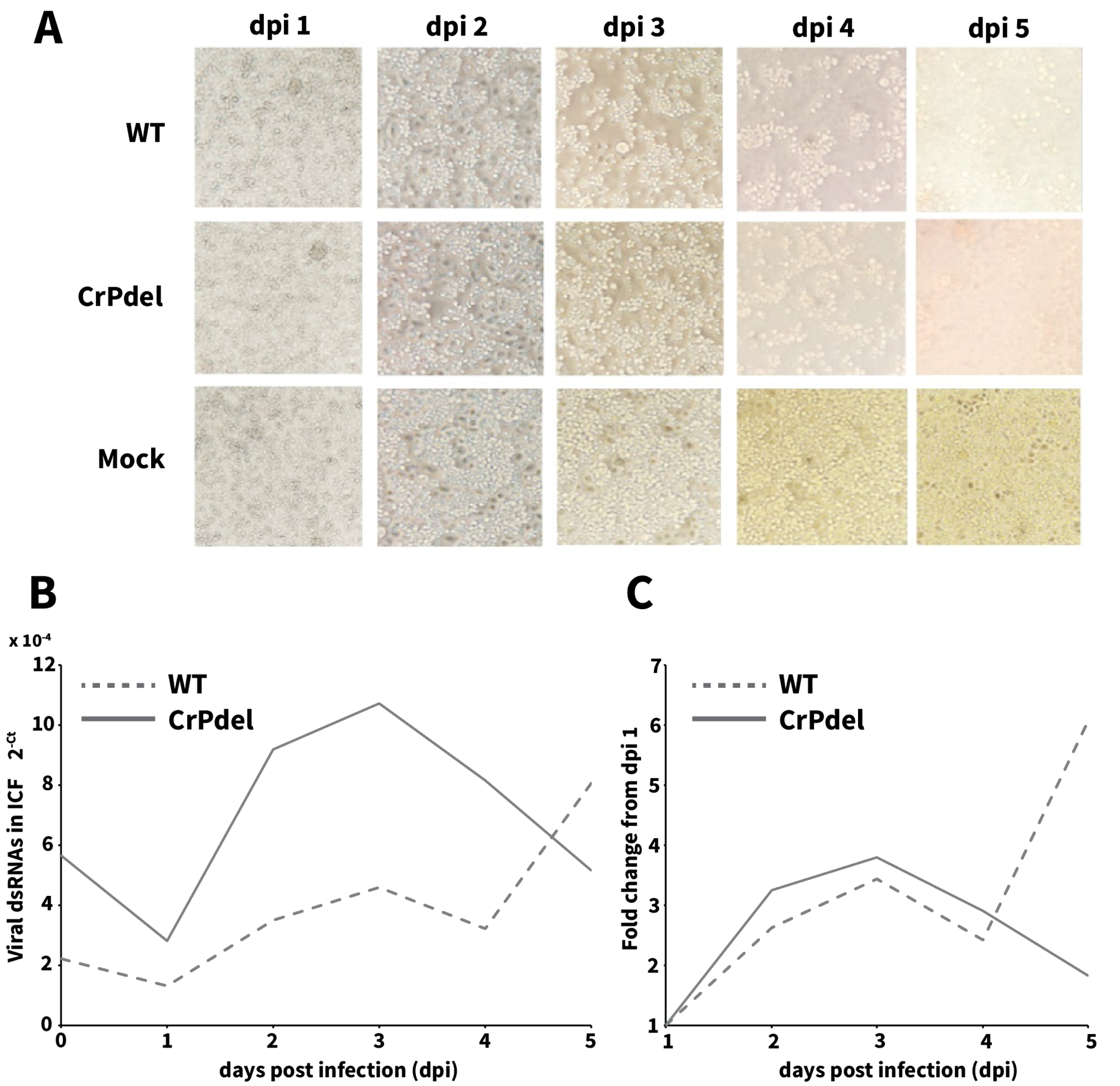


**Supplementary Fig. S1. CPE and propagation curves of OmRV-WT and OmRV-CrPdel after lipofection of the viral RNAs. (A)** CPE in C6/36 cells at 1, 2, 3, 4 and 5 dpi. C6/36 cells were introduced with OmRV-WT or OmRV-CrPdel viral RNAs (2.5 μg each). Mock indicates uninfected control cells. **(B)** Propagation efficiency of OmRV-WT and OmRV-CrPdel at 0, 1, 2, 3, 4, and 5 dpi following transfection with in vitro transcribed viral RNAs. **(C)** Fold change in viral RNA levels relative to those at 1 dpi.

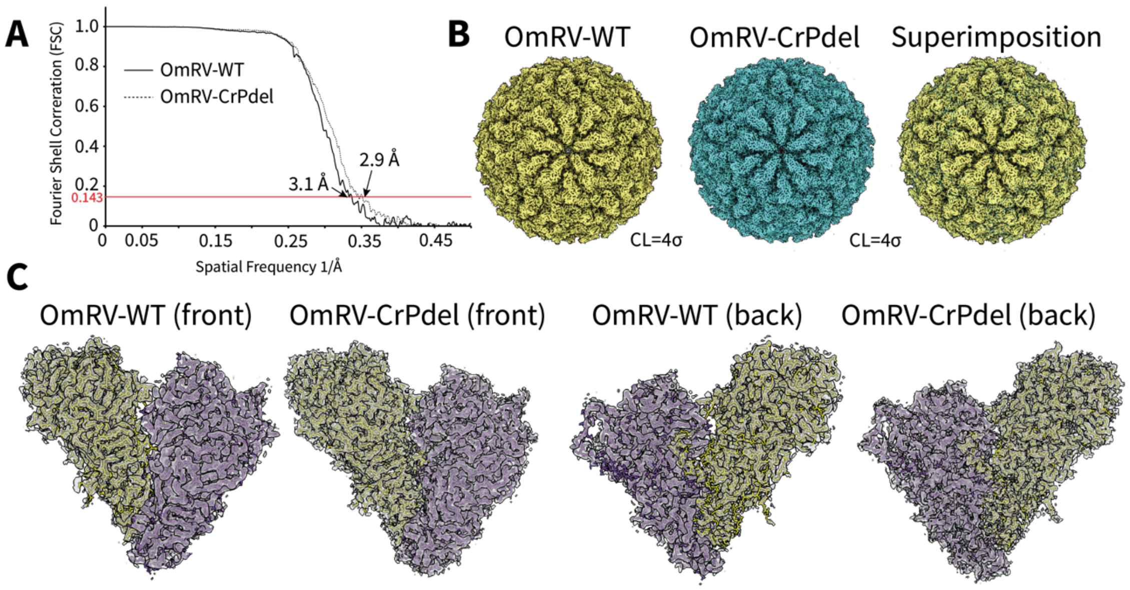


**Supplementary Fig. S2. Cryo-EM structural determination of OmRV-WT and OmRV-CrPdel. (A)** Fourier shell correlation (FSC) curves of the final OmRV-WT and OmRV-CrPdel reconstructions. **(B)** Overall structures of OmRV-WT and OmRV-CrPdel, and their superimposition. **(C)** Atomic model fitting for OmRV-WT and OmRV-CrPdel. The fitted models are colored by subunit, and the corresponding cryo-EM maps are shown as gray transparent surfaces.
